# Tomato cystatin SlCYS8 as a trigger of drought tolerance and tuber yield in potato

**DOI:** 10.1101/2024.09.11.612519

**Authors:** Maude Dorval, Andréane Langlois, Jonathan Tremblay, Gabrielle Veillet, Marc-Antoine Chiasson, Marie-Claire Goulet, Thiago Gumiere, Steeve Pepin, Charles Goulet, Dominique Michaud

**Affiliations:** Centre de recherche et d’innovation sur les végétaux, Université Laval, Québec QC, Canada; Département de phytologie, Université Laval, Québec QC, Canada; Département des sols et de génie agroalimentaire, Université Laval, Québec QC, Canada

**Keywords:** Potato (*Solanum tuberosum* L.), tomato cystatin SlCYS8, drought tolerance, tuber yield

## Abstract

Current climate change scenarios predict an increased incidence of drought episodes likely to affect potato crops worldwide. Potato exhibits a low-density, shallow root system that makes it particularly vulnerable to water shortage and any successful attempt to implement drought tolerance in cultivated potato varieties is potentially relevant from an agronomic standpoint. In this study, we assessed the potential of tomato cystatin SlCYS8 to promote drought tolerance in SlCYS8-expressing potato lines by induction of stress-related pleiotropy. Up to now, protease inhibitors of the cystatin protein superfamily have been mostly considered as biotechnological tools to engineer pest or pathogen resistance in crops, but several recent studies have also revealed a possible link between abiotic stress tolerance and these regulatory proteins in leaf tissue. SlCYS8-expressing plantlets grown on culture medium containing the drought mimic polyethylene glycol (PEG) exhibited an elevated root-to-shoot ratio, an indicator of drought tolerance in potato. A similar conclusion could be drawn with greenhouse-grown acclimated plants, confirming a relative root growth-promoting effect for the recombinant inhibitor upon water deficit. SlCYS8-potato lines also showed a high tuber yield compared to the control line under both limiting and non-limiting water regimes, suggesting an improved efficiency of the primary metabolism and the avoidance of a growth– stress response tradeoff in the modified lines. Accordingly, SlCYS8 expression was associated with a stress response-oriented proteome in leaves likely explained by pleiotropic effects of the recombinant cystatin driving the constitutive expression of stress-related proteins and the upregulation of primary metabolism-associated proteins. Overall, these data suggest the potential of cystatins as molecular triggers of tuber biomass production and drought resilience in potato. Complementary studies will be warranted to assess tuber yield of the SlCYS8-lines under different water regimes in field conditions.

## Introduction

Potatoes are an important source of nutrients for humans, with more than 350 million tons of tubers produced worldwide each year (FAO, 2024). At the global scale, potato ranks no. 1 among non-cereal food crops (Dahal et al., 2019) and no. 6 among major crops after sugarcane, corn, wheat, rice and palm oil (FAO, 2022). This crop grown on various soil types is now considered as a key player in the fight against hunger and malnutrition by the Food and Agriculture Organization (FAO, 2009). Potato tubers present a high nutritional value and provide an array of essential nutrients including carbohydrates, proteins, amino acids, fibers, vitamins and minerals (Beals, 2019). From a socioeconomic standpoint, farming and trading potatoes at the local scale promote food and economic security for human families and populations in many developing countries (FAO, 2009).

Despite interesting attributes, potato as a global source of energy and nutrients presents obvious challenges in the current context of a changing climate. This plant exhibits a low-density, shallow root system that makes it particularly susceptible to rising temperatures and a growing number of drought episodes every year (Dahal et al., 2019; Hill et al., 2021). Predictive models estimated that a temperature increase of 1 to 2 °°C would lead to an overall yield loss of 20%, and an increase of 2 to 3 °°C an overall loss of 30%, at the world scale (Hijmans, 2003). Modern potato varieties are ill prepared for future water deficits compared to their native counterparts bearing drought tolerance traits (Monneveux et al., 2013). Unfortunately, wild potato cultivars and Andean landraces are predominantly adapted to low latitude climates and are less suitable for farming on a global scale. Breeding efforts to transfer tolerance genes from native potato varieties to modern cultivars is an option for drought tolerance introgression, but this often results in undesirable outcomes such as a lower tuber yield or a high content of toxic alkaloids in tuber flesh (Monneveux et al., 2013).

Alternative strategies to classical breeding have been proposed for potato improvement, involving transgene addition by genetic transformation or specific, targeted alterations of the genome sequence by gene editing approaches (Hofvander et al., 2022; del Mar Martinez-Prada et al., 2021). In this study, we assessed the potential of plant cystatins as recombinant triggers of drought tolerance in cultivated potato. Members of the cystatin protein superfamily behave as pseudosubstrates to inhibit peptide bond hydrolysis in the active site cleft of cysteine (Cys) proteases (Benchabane et al., 2010; Martínez et al., 2012). In plants, cystatins and their Cys protease targets are involved in the regulation of key physiological processes including programmed cell death, protein deposition in storage organs and amino acid recycling in senescent leaves (Benchabane et al., 2010). Cystatins are also induced in leaves under various biotic or abiotic stress conditions and their practical potential in plant protection against herbivorous pests and pathogens has been discussed extensively over the past decades (reviewed in Tremblay et al., 2019). Meanwhile, research efforts have been deployed to explore the usefulness of these proteins in alleviating abiotic stress-induced symptoms under adverse cultural or environmental conditions like soil salinity, soil alkalinity, drought and suboptimal temperatures (Van der Vyver et al., 2003; Prins et al., 2008; Zhang et al., 2008; Quain et al., 2014; Je et al., 2014; Sun et al., 2014; Tan et al., 2015, 2017; Liu et al., 2020).

At this point, the mode of action of inhibitory cystatins is well characterized at the biochemical scale but the physiological effects of plant cystatins remain elusive for the most part. A useful, generic model to explain the action of these proteins in plants is the establishment of a high cystatin–Cys protease stoichiometric ratio upon cystatin induction or heterologous expression, which in turn influences Cys protease-dependent physiological functions *in planta* (Benchabane et al., 2010). In potato, the hypothesis of a stoichiometric ratio-driven impact for cystatins is well supported by an elevated multicystatin–Cys protease ratio in growing tubers accumulating storage proteins (Weeda et al., 2009) and, conversely, a low cystatin–Cys protease ratio in germinating tubers hydrolyzing storage proteins for amino acid mobilization (Weeda et al., 2010). The physiological impacts of a high cystatin content in potato were also illustrated by reduced Cys protease activities leading to delayed sprouting in stored tubers expressing rice cystatin I (Munger et al., 2015), or by the induction of stress proteins and Cys protease-independent resistance to the necrotrophic pathogen *Botrytis cinerea* in potato lines engineered to express corn cystatin II (Munger et al., 2012). Here, we assessed the potential of SlCYS8, a model cystatin from tomato (Goulet et al., 2008), to promote drought tolerance in potato via the induction of stress-related pleiotropic effects in leaf tissue.

## Results and Discussion

### SlCYS8-potato lines present a stress protein-enriched leaf proteome

Transgenic potato lines expressing tomato SlCYS8 and the antibiotic selection marker neomycin phosphotransferase II were regenerated *in vitro* on kanamycin-containing medium following *Agrobacterium*-mediated transformation with a synthetic version of the SlCYS8 DNA coding sequence (**Fig. 1A**; **Supplementary Fig. S1**). *In vitro* clones obtained from independent calli were acclimated in the greenhouse and PCR-tested for the selection marker gene using appropriate DNA primers. A ∼500-base-long ‘nptii’ amplicon was amplified from the genomic DNA of all tested plants, confirming that all clones regenerated on kanamycin-medium had been genetically transformed by the bacterial vector. Immunodetections were performed with rabbit anti-SlCYS8 polyclonal IgG to compare SlCYS8 levels in leaves among the regenerated clones. As expected, given the random site insertion of transgenes in the genome of *Agrobacterium*-infected cells, the immunoblot signals for SlCYS8 at ∼10 kDa differed from one clone to another, from null [undetectable] (e.g., clones C54, C55 and C57) or very weak (clones C44 and C53) to moderate (clones C3, C6, C56 and C58) or strong (clones C41, C43 and C51) (**Fig. 1B**). A liquid chromatography-tandem mass spectrometry (LC-MS/MS) analysis was performed on leaf protein extracts of control line K and transgenic clones C41, C43 and C51 to confirm an increased amount of cystatin peptides in the SlCYS8-expressing clones (**Fig. 1C**). Potato leaves naturally produce a stress-inducible, insoluble 88-kDa multidomain cystatin, potato multicystatin, that can be found at some level in healthy leaves (Munger et al., 2012). Accordingly, cystatin peptides presumably derived from the endogenous inhibitor domains were detected in leaves of control line K. By comparison, cystatin peptide intensities in lines C41, C43 and C51 were two to three times greater, in line with SlCYS8 heterologous expression in the modified lines likely leading to an increased cystatin–Cys protease stoichiometric ratio in leaf tissue.

**Figure 1.**
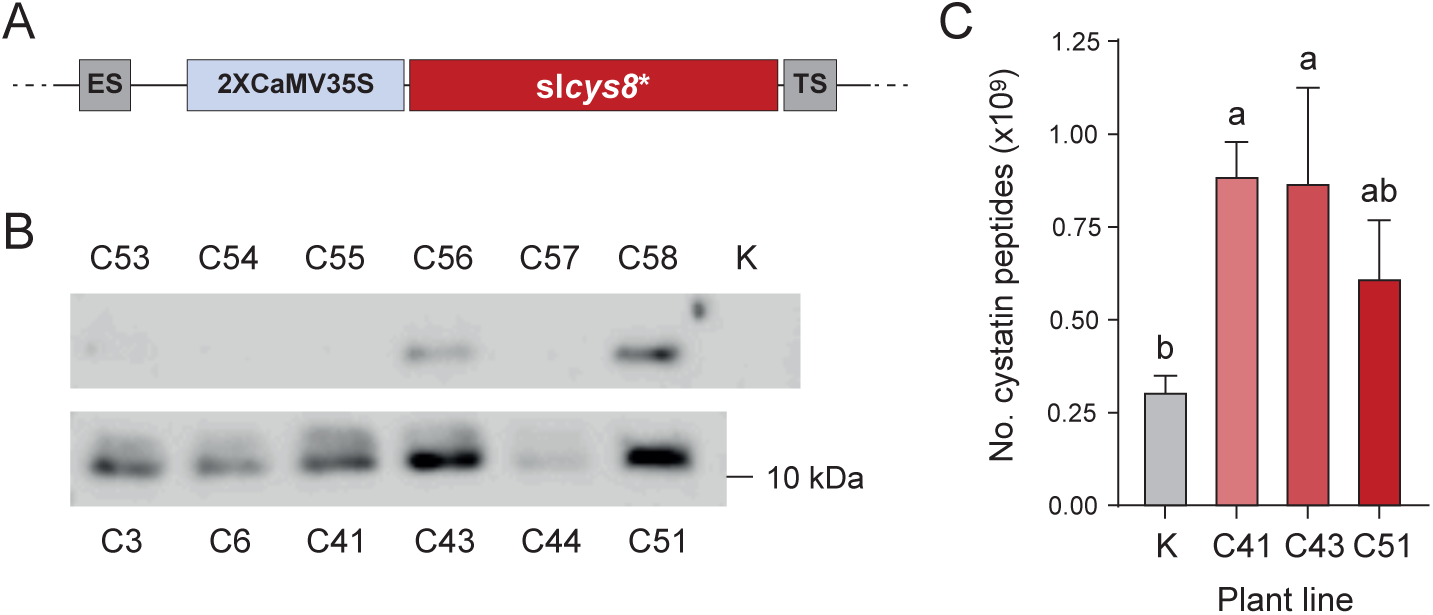
SlCYS8 heterologous expression in potato. (**A**) Gene construct elements for the cytosol-targeted expression of SlCYS8 in potato cells. The construct included an optimized version of the slcys8 sequence (slcys8*; see **Supplementary Fig. S1**), a duplicated version of the Cauliflower mosaic virus 35S (2xCaMV35S) promoter for constitutive expression, a tobacco etch virus enhancer sequence (ES) in upstream position of the cystatin sequence and a CaMV35S terminator sequence (TS) in downstream position. (**B**) Immunoblot signals for recombinant SlCYS8 in selected kanamycin-resistant lines. Immunodetections were performed using polyclonal IgG raised in rabbit against the tomato cystatin. (**C**) Cystatin peptide numbers in control (parental) line K and SlCYS8-expressing lines C41, C43 and C51, as determined following nanoLC-MS/MS. Each bar is the mean of 4 biological replicate values ± SD. Bars with different letters are significantly different (post-ANOVA Tukey’s test; α = 0.05)

We conducted a detailed analysis of our LC-MS/MS dataset to assess the possible impacts of SlCYS8 expression on the host plant stress proteome. Several studies have reported endogenous protein alterations in plants expressing a recombinant cystatin (Tremblay et al., 2019). In potato, heterologous expression of corn cystatin II (CCII) was shown to upregulate normally inducible biotic stress-related (defense) proteins in leaves, including pathogenesis-related (PR) protein PR-1, PR-2 proteins (ß-glucanases), PR-3 proteins (chitinases) and protease inhibitors (Munger et al., 2012). CCII heterologous expression was also associated with the upregulation of abiotic stress-related proteins involved in osmotic and oxidative stress mitigation, such as osmotin and secretory peroxidases (Munger et al., 2012). Supporting these previous observations and confirming the successful expression of tomato SlCYS8 under an active form in potato cells, numerous alterations of the leaf proteome were noted in lines C41, C43 and C51 compared to control line K. Out of 3,141 proteins detected overall (**Supplementary Dataset S1**), 175 proteins were differentially regulated in the transgenic lines, including 122 upregulated proteins and 53 downregulated proteins (**Fig. 2**). A Gene Ontology (GO) enrichment analysis was conducted for those 151 proteins among the 175 differentially expressed proteins that were significantly regulated in at least one, and detected in at least two, of the three tested lines. Of these 151 proteins, 53 were identified as being likely associated with biotic stress (defense) or abiotic stress responses (**Fig. 3**; **Supplementary Table S1**).

**Figure 2.**
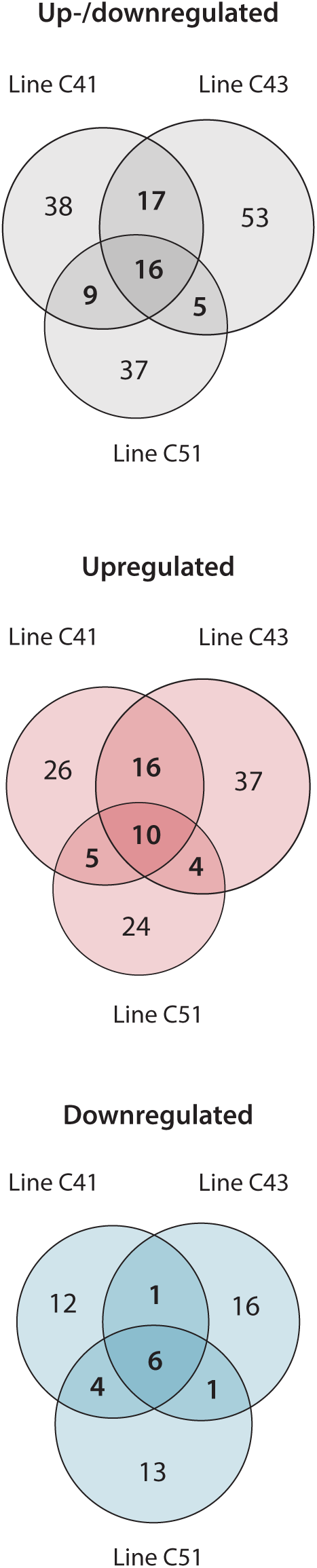
Venn diagrams for up- and/or downregulated proteins in leaf tissue of SlYCS8-expressing lines C41, C43 and C51. Control (parent) line K was used as a baseline for the comparisons. Numbers in overlapping areas indicate numbers of proteins regulated in two or all three transgenic lines compared to the control line, out of 175 regulated proteins overall (|z-score| ≥ 1.96l *p*-value < 0.05) (see **Supplementary Dataset S1**).

**Figure 3.**
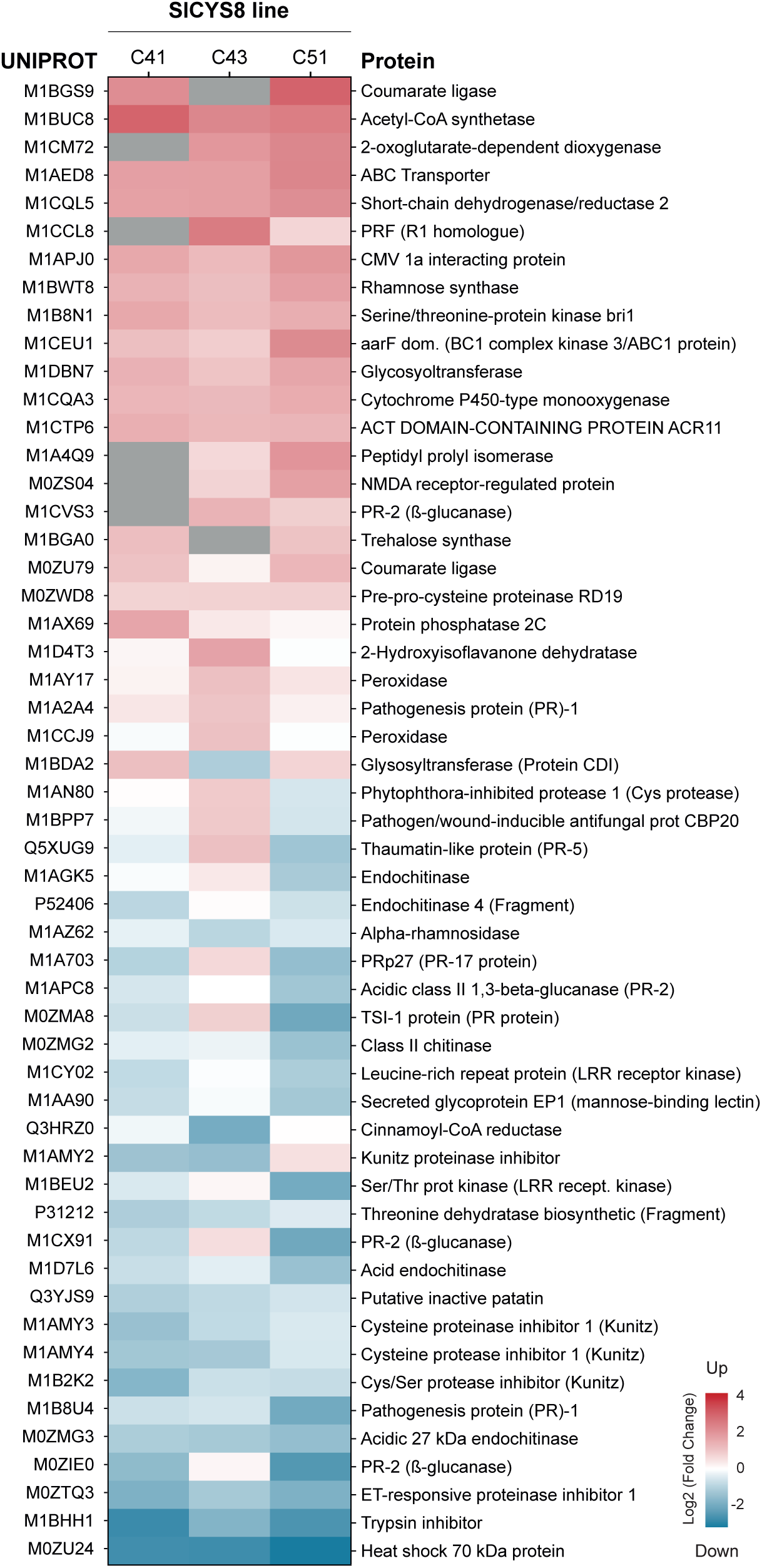
Defense/stress-related proteins up- or downregulated in SlCYS8-expressing lines C41, C43 and C51. Proteins on this figure were significantly up- or downregulated in at least one, and detected in at least two, of the three tested lines. Upregulated proteins are shown in red, downregulated proteins in blue. A grey area indicates no detection of the protein. Log2 values of peptide ratios for each line compared to parent line K are provided in Supplementary Table S1.

Upregulated defense-related proteins included several salicylic acid-inducible PR proteins, protease inhibitors and key enzymes of the phytoalexin and coumarin biosynthetic pathways including acetyl-CoA synthetase, coumarate ligase, 2-oxoglutarate-dependent dioxygenase and cinnamoyl-CoA reductase. Proteins known to alleviate the detrimental effects of abiotic stresses were also found in the upregulated fraction (**Fig. 3**), including xenobiotics-metabolizing enzymes and oxidative stress-mitigating enzymes such as secretory peroxidases, a cytochrome P450 monooxygenase and several enzymes of the flavonoid biosynthetic pathway (Kidwai et al., 2020; Pandian et al., 2020; Shomali et al., 2022). An ATP-binding cassette (ABC) transporter and a peptidyl-prolyl isomerase catalyzing proline biosynthesis were also upregulated, in line with the well-documented protective functions of proline and ABC transporters in adverse conditions (Dahuja et al., 2021; Ghosh et al., 2021). These observations suggesting the onset of a stress protein-enriched proteome in the SlCYS8-potato lines were in agreement with previous studies reporting increased antioxidant enzyme activities and reduced accumulation of reactive oxygen species, including hydrogen peroxide, in Arabidopsis, tobacco and apple engineered to express a recombinant cystatin (Demirevska et al., 2010; Tan et al., 2017, 2016, 2015).

Unexpectedly, several defense proteins were downregulated in the SlCYS8-lines, including protease inhibitors, acidic endochitinases and ß-glucanases naturally induced by the defense elicitor jasmonic acid (**Fig. 3**). This observation, together with the upregulation of salicylic acid-inducible proteins in the modified lines (see above), prompted us to look for a possible impact of SlCYS8 expression on the relative concentrations of salicylic acid and jasmonic acid in leaf tissue. These two defense elicitors present an antagonistic relationship *in planta* and it is well established that an increase in salicylic acid cellular content generally compromises the expression of jasmonic acid-inducible genes (Van der Does et al., 2013). To address this, an ultraperformance liquid chromatography (UPLC)-MS/MS procedure was followed to compare the hormone profiles of SlCYS8-expressing lines C41, C43 and C51 with the hormone profile of control line K (**Supplementary Fig. S2**). In brief, the basic levels of auxin, cytokinin, abscisic acid and 1-aminocyclopropane-1-carboxylic acid (ACC), the precursor of ethylene, were roughly comparable in the four tested lines, suggesting a limited impact of SlCYS8 expression on the content of these hormones in leaf tissue. By comparison, jasmonic acid content was slightly lower in the SlCYS8-expressing lines, possibly explaining in part the observed downregulation of jasmonic acid-inducible proteins in leaves. Salicylic acid and its conjugated forms were also found at slightly lower levels in the SlCYS8-potato lines, eventually ruling out the possibility of a salicylic acid-dependent repression of the jasmonate-inducible proteins, and the possibility of a salicylic acid-dependent upregulation of salicylic acid-inducible proteins upon SlCYS8 expression. Together, these observations suggested the onset of hormone-independent pleiotropic effects for the recombinant cystatin likely involving the sequential inhibition of (a) [still to be identified] Cys protease target(s) in leaves and a subsequent hormone-independent up- or downregulation of several components of the stress/defense-related leaf proteome.

### SlCYS8-potato and control K lines show contrasting phenotypes under well-watered and water deficit treatments

Our MS/MS data showing the onset of a stress protein-enriched proteome in the SlCYS8-expressing lines supported the idea of an eventual beneficial impact for the recombinant cystatin under water deficit conditions. This assumption was also suggested by previous transcriptomic studies highlighting the preferential expression of stress-related proteins in drought-tolerant potato genotypes (Watkinson et al., 2006; Aliche et al., 2022). Stress proteins in these lines included oxidation-mitigating enzymes, cytochromes P450 and different enzymes of the flavonoid and phenylpropanoid biosynthetic pathways, similar to the population of stress proteins here identified as upregulated in our proteome dataset for the SlCYS8-expressing lines.

A water deficit assay was performed with *in vitro*-grown potato plantlets as a first step to assess the drought tolerance status of lines C41, C43 and C51, using the osmotic stress agent polyethylene glycol (PEG) as a mimic for water stress (Anithakumari et al., 2011) (**Fig. 4**, **Supplementary Table S2**). As expected, given the well-documented negative impact of water deficit on leaf cell growth, increasing concentrations of PEG in the growth medium, from 0 to 40 mM, induced a gradual decline of plantlet height and shoot (aboveground) biomass, easily observed after 4 weeks for all tested lines (*see* **Fig. 4A** for control line K). Despite a reduced growth rate, SlCYC8-expressing plantlets grown on 20 mM PEG medium exhibited a substantially increased root-*to*-shoot dry biomass ratio (**Fig. 4B**), a positive marker of drought tolerance in potato (Schafleitner et al., 2007). Variable, non-conclusive effects were observed at 30 mM and 40 mM PEG for lines C43 and C51 (not shown), likely indicative of severe stress effects above the host plant’s ability to tolerate the osmotic agent stressor. By comparison, control line K showed a low root-*to*-shoot biomass ratio at 20 mM compared to the SlCYS8-potato lines, roughly comparable to its root-*to*-shoot biomass ratio at 0 mM, 30 mM or 40 mM PEG.

**Figure 4.**
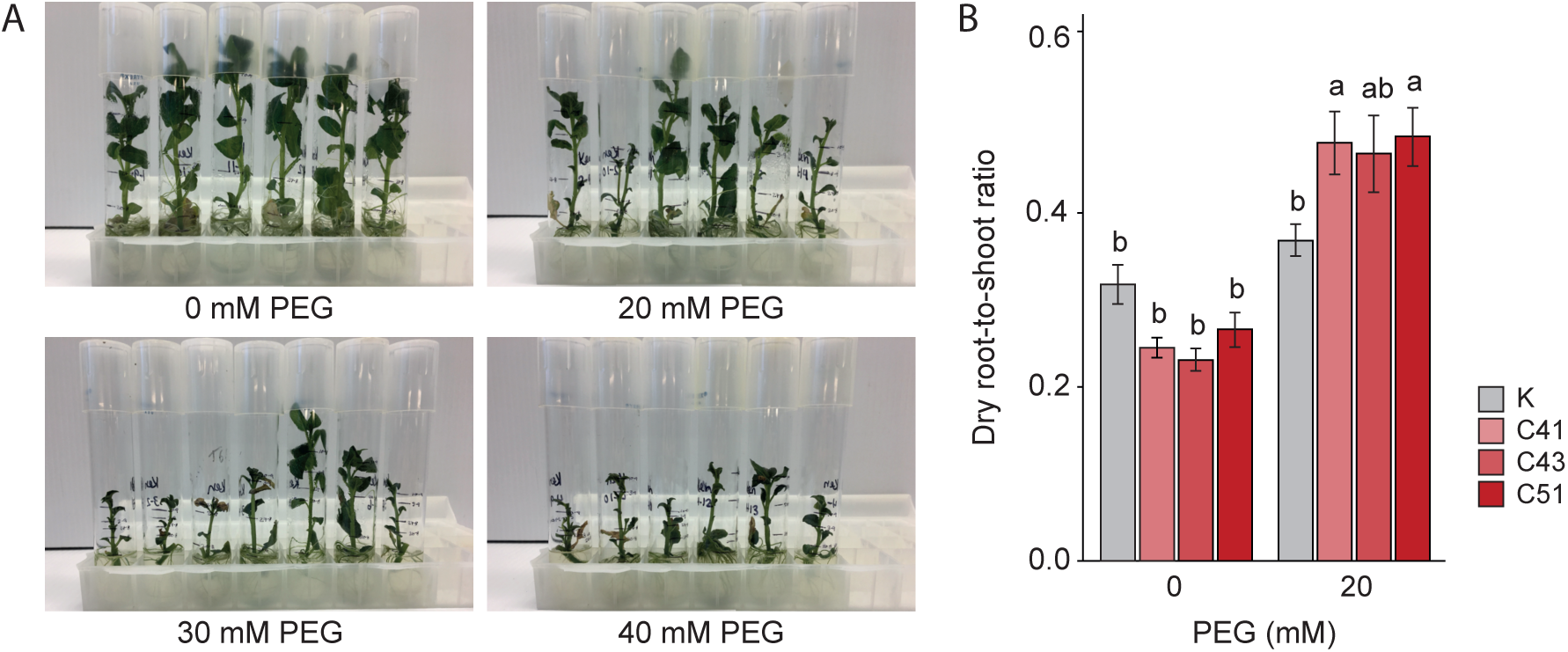
Growth and dry root-to-dry shoot ratio of parent line K and SlCYS8-expressing lines C41, C43 and C51 grown *in vitro* under increasing PEG concentrations. (**A**) Visual assessment of line K *in vitro* plantlets grown for 28 d under increasing PEG concentrations. (**B**) Dry root-*to*-shoot weight ratio of control and SlCYS8-expressing plantlets grown for 28 d under 0 mM (no-stress control) or 20 mM PEG. Each bar on panel B is the mean of 4 biological replicate values ± SEM. Bars with different letters at a given PEG concentration indicate significantly different mean values (post-ANOVA Tukey’s test; α = 0.05).

A complementary experiment was conducted in greenhouse with fully-grown, acclimated plants to confirm the apparent ability of the SlCYS8-potato lines to better sustain the relative biomass of their root system on a whole plant basis and to assess the ability of these lines to grow and produce tubers under water deficit conditions over a complete production cycle. Acclimated plantlets were placed in 0.14 m^3^ units filled with a handmade blend of sand and organic soil assembled to present good drainage and a propensity to generate water deficit conditions. The plants were grown to full maturity (i.e., up to the flowering stage) under well-irrigated conditions. Three water treatments were then applied until tuber harvest, in such a way as to generate well-watered (–5 kPa) and water deficit (–10 kPa, –15 kPa) conditions. The substrate matric potential was monitored without interruption to ensure stable water conditions throughout the experiment. Several above- and belowground measurements were taken at harvest, as summarized in **Table 1**.

**Table 1.** Growth parameters of greenhouse-grown control and SlCYS8-expressing potato lines under well-irrigated or water stress regimes. Data were collected after 90 d in greenhouse, i.e. 40 d after initiating the water stress treatments. Values are the mean of 4 biological replicate values ± SEM. Refer to **Supplementary Table S3** for additional details on statistical inferences.

| Irrigation threshold (kPa) | Potato line | Stem length (cm) | Stem number | Fresh shoot biomass (g) | Dry shoot biomass (g) | Root length (cm) | Dry root biomass (g) | Dry root-to-shoot ratio |
| --- | --- | --- | --- | --- | --- | --- | --- | --- |
| – 5 | K | 103.0 ± 2.6 | 5.1 ± 0.4 | 369 ± 14 | 38.2 ± 1.5 | 57.5 ± 6.1 | 1.6 ± 0.08 | 4.6 ± 0.39 |
| | C41 | 82.4 ± 8.2 | 6.9 ± 0.3 | 327 ± 17 | 30.7 ± 2.0 | 46.4 ± 4.4 | 1.4 ± 0.16 | 4.7 ± 0.45 |
| | C43 | 89.1 ± 5.0 | 8.1 ± 1.2 | 366 ± 15 | 35.9 ± 1.0 | 52.4 ± 4.2 | 1.5 ± 0.08 | 4.3 ± 0.32 |
| | C51 | 86.6 ± 4.2 | 6.9 ± 0.7 | 350 ± 28 | 32.1 ± 2.5 | 56.9 ± 9.3 | 1.6 ± 0.07 | 5.1 ± 0.40 |
| – 10 | K | 96.8 ± 6.8 | 4.8 ± 0.1 | 317 ± 18 | 34.4 ± 1.0 | 45.1 ± 4.0 | 1.5 ± 0.12 | 4.3 ± 0.25 |
| | C41 | 84.0 ± 5.6 | 5.6 ± 0.6 | 316 ± 22 | 32.4 ± 1.7 | 47.4 ± 3.1 | 1.3 ± 0.06 | 4.2 ± 0.09 |
| | C43 | 85.6 ± 3.0 | 6.8 ± 0.3 | 319 ± 22 | 29.8 ± 3.3 | 48.4 ± 3.6 | 1.4 ± 0.10 | 4.9 ± 0.22 |
| | C51 | 92.1 ± 5.9 | 7.1 ± 0.4 | 331 ± 28 | 32.5 ± 2.5 | 57.3 ± 2.6 | 1.5 ± 0.14 | 4.7 ± 0.38 |
| – 15 | K | 89.6 ± 4.8 | 5.6 ± 0.4 | 256 ± 12 | 27.2 ± 1.0 | 45.4 ± 3.0 | 1.3 ± 0.06 | 4.7 ± 0.24 |
| | C41 | 73.8 ± 2.2 | 7.8 ± 0.3 | 256 ± 10 | 25.1 ± 0.4 | 47.3 ± 2.7 | 1.4 ± 0.06 | 5.6 ± 0.32 |
| | C43 | 74.6 ± 3.9 | 7.6 ± 0.2 | 231 ± 20 | 23.2 ± 1.0 | 49.8 ± 2.4 | 1.4 ± 0.04 | 6.3 ± 0.17 |
| | C51 | 78.6 ± 3.2 | 7.5 ± 0.7 | 252 ± 8 | 23.7 ± 0.7 | 44.6 ± 4.2 | 1.4 ± 0.04 | 6.0 ± 0.34 |
| Block |  | ** | * | n.s. | n.s. | *** | ** | * |
| Threshold |  | ** | * | *** | *** | n.s. | * | *** |
| Potato line |  | *** | *** | n.s. | * | n.s. | n.s. | * |
| Threshold x Potato line |  | n.s. | n.s. | n.s. | n.s. | n.s. | n.s. | n.s. |

\*\*\*, $P < 0.001$ ; \*\*, $P < 0.01$ ; \*, $P < 0.05$ ; n.s., not significant (two-way ANOVA; $\alpha = 0.05$ ).

Significant differences were observed among treatments after 90 d for most of the traits considered, thereby confirming the validity of our water deficit treatments. Transgenic lines differed from the control line in terms of aboveground biomass distribution, with the control plants presenting longer stems and SlCYS8-expressing lines harbouring a larger number of shoots (**Table 1**). In a previous study, Quain et al. (2014) reported a shoot branching phenotype for soybean lines expressing rice cystatin I, reminiscent of the shoot branching phenotype of an Arabidopsis mutant for CCD7 and CCD8, two key enzymes of the strigolactone biosynthetic pathway. Here, the underlying cause of an increased shoot branching in the SlCYS8-expressing lines remains to be determined. A physiological effect implicating strigolactone deficiency upon SlCYS8 heterologous expression cannot be ruled out at this stage but this scenario appears unlikely considering the above-described leaf hormonal profiles of control line K and SlCYS8-expressing lines C41, C43 and C51 (**Supplementary Fig. S2**). The net impact of strigolactones on shoot branching is generally explained by a positive effect on auxin biosynthesis and/or a concomitant negative effect on cytokinin biosynthesis (Barbier et al., 2019; Wang et al., 2019). Based on current models, any change in shoot branching pattern implicating strigolactones would be associated with an altered auxin/cytokinin balance *in planta* that does not match the unchanged concentrations of auxins and cytokinins observed above in the SlCYS8-expressing lines.

As expected, given the well-described negative impact of water deficit on plant cell growth, plant canopy biomass decreased under water stress for both the SlCYS8-expressing and control lines (**Table 1**). More specifically, the four lines showed a similar aboveground fresh biomass for both the well-irrigated and water deficit treatments. Minor differences were observed among the lines for the dry aboveground biomass but the differences were marginal, as indicated by a non-significant post-hoc Tukey test on the average biomass values (**Supplementary Table S3**). As for the aboveground biomass, dry root biomass decreased with water deficit intensity, but the length of the root system remained unchanged among the lines under the different water regimes (**Table 1**). By contrast, and as observed above with the *in vitro* plantlets, the SlCYS8-expressing lines exhibited an increased root-*to*-shoot biomass ratio under water deficit, greater overall than the root-*to*-shoot ratio determined for the control line, especially for the most pronounced water deficit treatment at –15 kPa. Considering the stress protein enrichment effects of SlCYS8 on the leaf proteome, the ability of the SlCYS8-expressing lines to sustain a greater root-*to*-shoot dry biomass ratio under water deficit, either *in vitro* and *ex vitro*, raised the possibility of an improved tuber yield for these lines compared to the control line, eventually associated with an improved cellular machinery for stress mitigation under adverse conditions.

### SlCYS8-potato lines produce higher tuber yields under both well-irrigated and water deficit treatments

In accordance with the negative impact of water deficit on plant canopy biomass, total tuber biomass was negatively correlated with water deficit intensity for all tested lines (**Fig. 5A**). The total number of tubers per plant remained unaltered upon water stress (**Fig. 5B**), thus suggesting no direct effect of SlCYS8 expression on tuber initiation. As expected, given the proteomic and phenotypic inferences above, total tuber biomass produced by the SlCYS8 lines was greater than the control line under water deficit treatments (**Fig. 5A**). Much interestingly, total tuber biomass was ∼50% greater for the SlCYS8 lines under the well-watered treatment, indeed confirming a positive impact of SlCYS8 expression not only on the stress response but also on basic, yield-associated primary metabolic functions.

**Figure 5.**
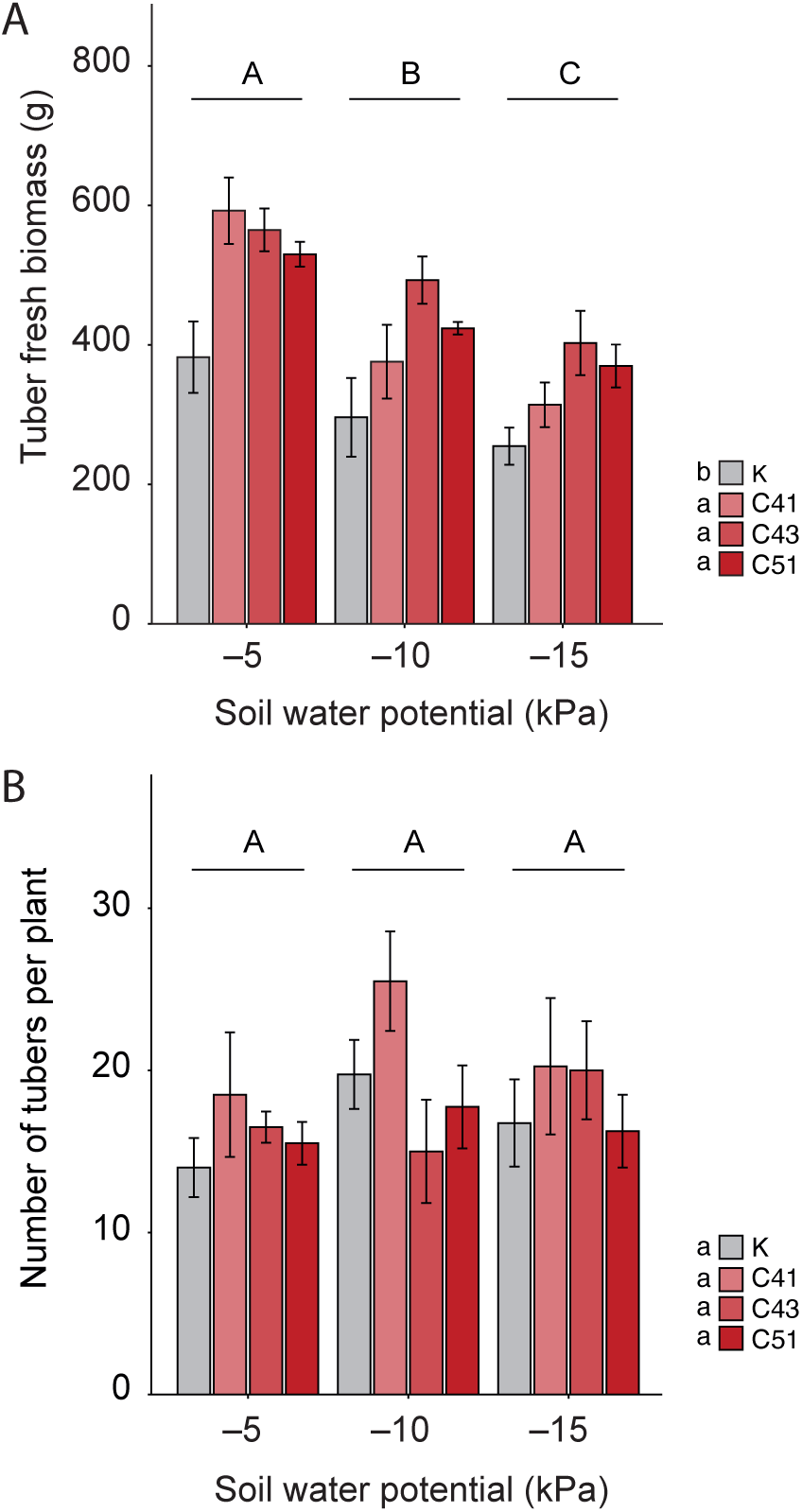
Tuber yield of parent line K and SlCYS8-expressing lines C41, C43 and C51 grown in greenhouse under well-watered or water stress conditions. (**A**) Total tuber biomass per plant. (**B**) Number of tubers per plant. Each bar on panel A or panel B is the mean of 4 biological replicates ± SEM. Different letters on a given panel indicate significant differences among the lines (lowercase letters) or water stress treatments (uppercase letters) (post two-way ANOVA Tukey’s test; α = 0.05).

We revisited our MS/MS data to look for leaf proteome alterations eventually supporting the idea of a strengthened primary metabolism in lines C41, C43 and C51 (**Fig. 6**; **Supplementary Table S4**). Of the 122 proteins identified above as upregulated in the SlCYS8-expressing lines compared to control line K (**Supplementary Dataset S1**), 63 proteins could be assigned to GO functional categories directly or indirectly related to cellular metabolism, plant growth and/or tuber yield. Thirty-two (32) proteins involved in protein biosynthesis and maturation, including several ribosomal proteins and eukaryotic translation factors, and 11 proteins involved in protein transport or trafficking were enriched in the transgenic lines (**Fig. 6A,B**), suggesting a general amplification of the protein biosynthetic machinery upon recombinant cystatin expression. Seven (7) starch-/sugar-metabolizing enzymes and 13 photosynthesis-associated proteins, including chlorophyll a/b-binding proteins, ferredoxin and thioredoxin, were also found at higher levels in the transgenic lines (**Fig. 6C,D**), in accordance with greater tuber biomass for the SlCYS8-expressing lines under non-limiting water conditions (**Fig. 5A**) and the previously reported preferential expression of starch- and carbohydrate-processing genes in drought-tolerant potato varieties (Aliche et al., 2022).

**Figure 6.**
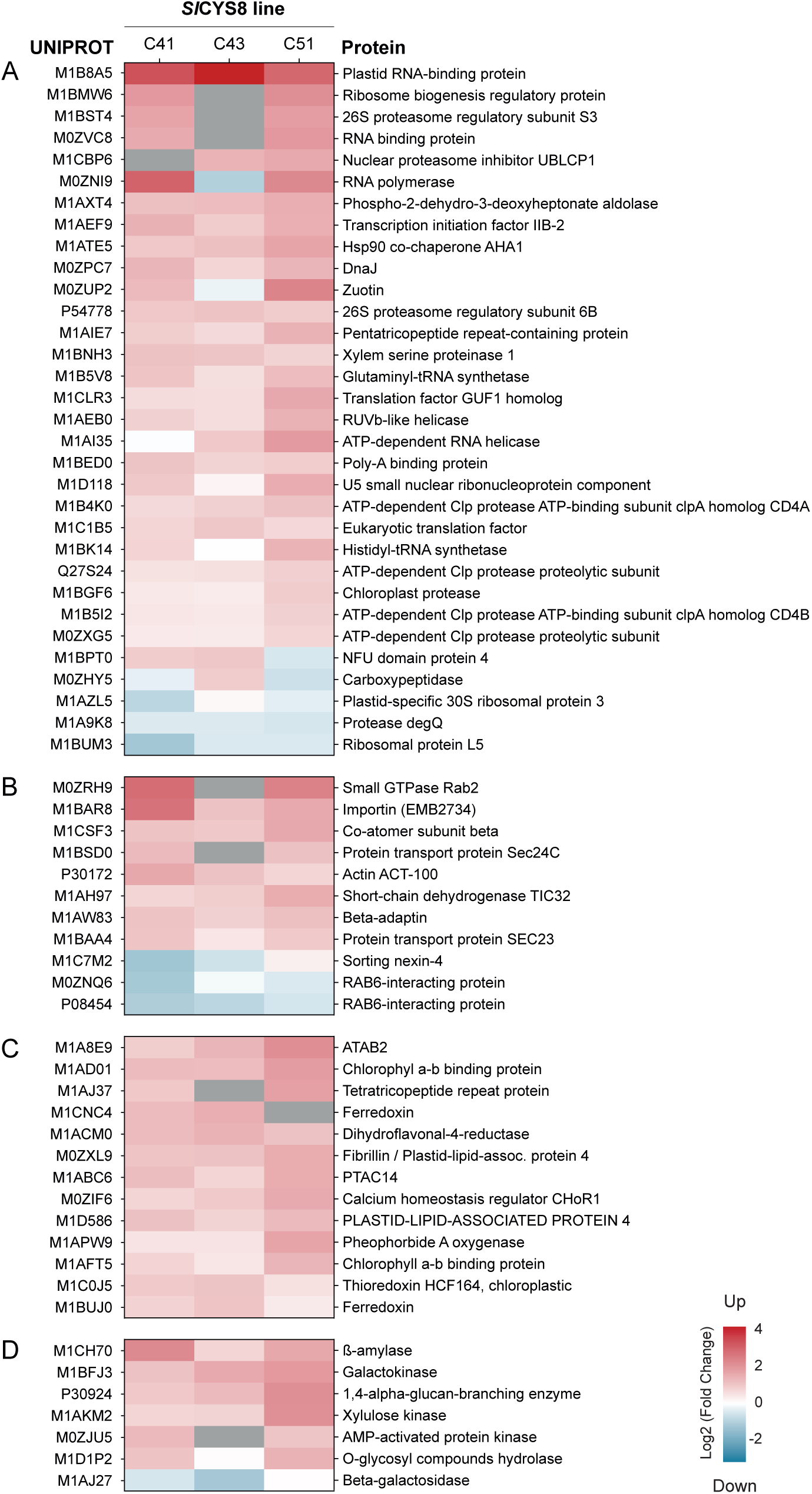
Primary metabolism-associated proteins up- or downregulated in SlCYS8-expressing lines C41, C43 and C51. (**A**) Proteins involved in protein biosynthesis, maturation or processing. (**B**) Proteins involved in protein transport or cellular trafficking. (**C**) Proteins involved in photosynthesis or photosystems assembly/maintenance. (**D**) Enzymes involved in carbon or sugar metabolism. Proteins on this figure were significantly up- or downregulated in at least one, and detected in at least two, of the three tested lines. Upregulated proteins are shown in red, downregulated proteins in blue. A grey area indicates no detection of the protein. Log2 values of peptide ratios for each line compared to parent line K are provided in Supplementary Table S4.

We calculated drought tolerance (DTI) and yield performance (YTI) indices to determine whether higher tuber yields for the SlCYS8 lines upon water deficit were associated with an actual drought tolerance phenotype or, alternatively, with an improved metabolic status sustaining higher yields in water-limiting conditions as a result of upstream yield-promoting pleiotropic effects already giving the plant an advantage in non-limiting conditions (**Fig. 7**). DTI measures the drought tolerance status of a given genotype, as defined in terms of yield preservation under limiting water conditions compared to the average DTI index of all considered genotypes, while YTI discriminates a given genotype based on its yield potential compared to the other genotypes under all tested conditions (Ober et al., 2004). In brief, no significant difference was observed between the lines or the irrigation treatments for the DTI indices, suggesting a general, drought-independent yield-promoting effect of the recombinant cystatin in lines C41, C43 and C51 under water deficit conditions. This conclusion was supported by greater YTI indices for the transgenic lines, estimated at about two times the YTI indices calculated for control line K under the water deficit treatments.

**Figure 7.**
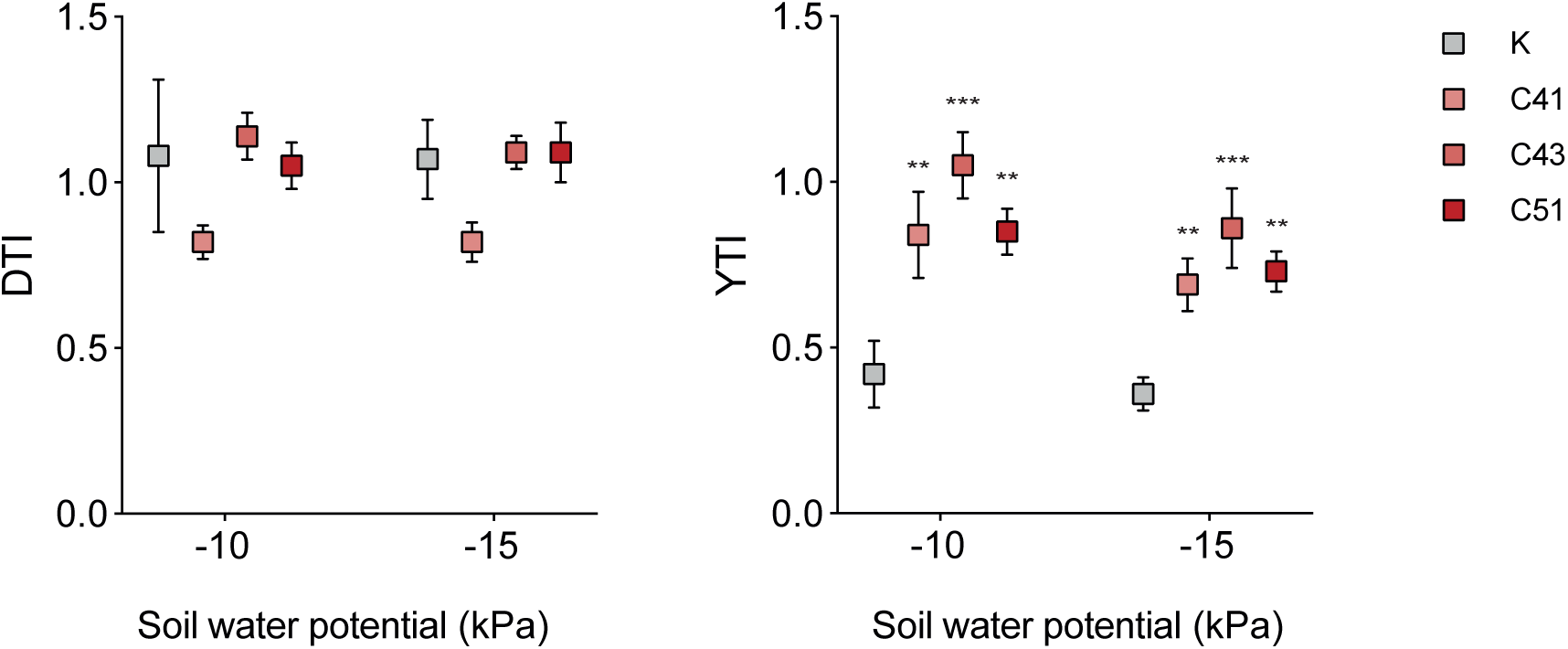
Drought tolerance indices (DTI) and Yield performance indices (YTI) of control line K and SlCYS8-expressing lines C41, C43 and C51 grown under water stress regimes. Data are compared to the no-stress water regime at –5 kPa. Values are the mean of 4 biological replicate values ± SEM. Asterisks indicate significantly different indices compared to control line K (two-way ANOVA; α = 0.05; **, *P* < 0.01, *** *P* < 0.001).

## Conclusion

Several papers over the last decade highlighted a possible link between plant abiotic stress tolerance (or resilience) and the occurrence of cystatins in leaf tissue. Our goal in this study was to assess the potential of a well-characterized plant cystatin, tomato SlCYS8 (Goulet et al., 2008), to trigger water deficit tolerance in cultivated potato by the induction of stress-related pleiotropic effects. Supporting this eventuality, transgenic potato lines expressing SlCYS8 presented a stress protein-enriched leaf proteome and a significantly increased root-*to*-shoot biomass ratio under water deficit conditions, either *in vitro* with non-acclimated plantlets and *ex vitro* after plant acclimation. Unexpectedly, given the usual growth–defense tradeoffs observed in plants coping with stressful environments (Claeys and Inzé, 2013) and the often poor agronomic performance of potato accessions presenting abiotic stress tolerance traits (Monneveux et al., 2013), SlCYS8 expression also had a positive impact on tuber yield, associated with the upregulation of growth-, photosynthesis- and yield-associated proteins in leaves. Overall, these observations confirmed the manifestation of multiple SlCYS8-induced pleiotropic effects in potato promoting its general performance under both favorable and less favorable water regimes.

Studies will be warranted in coming years to validate the observed positive effects of SlCYS8 on tuber yield and drought tolerance (or resilience) in field conditions, and to assess the potential of alternative cystatin candidates for the improvement of potato cultivars presenting different degrees of drought tolerance. Studies will also be welcome to further characterize the *in planta* physiological effects of SlCSY8, from the identification of its putative protease target(s) to the elucidation of the cellular and/or gene regulatory processes driving the observed alterations of the leaf proteome. Work is now planned to identify the protease target(s) of SlCYS8 in potato leaves, as a first step toward the design of a genome editing alternative to the SlCYS8 transgene, ‘GM-based’ strategy relying on post-expression protease inhibition.

## Materials and methods

### SlCYS8 transgene construct

Transgenic potato lines expressing a codon-optimized version of the SlCYS8 DNA sequence were used for the experiments to prevent silencing in leaves expressing potato multicystatin, a close homologue of the tomato cystatin. The codon-optimized sequence, *slcys8\**, was designed *in silico* by the inclusion of synonymous mutations, in such a way as to preserve the original amino acid sequence (**Supplementary Fig. S1**). A DNA template for the synthetic transgene was synthesized as a DNA g-block™ (Integrated DNA Technologies, inc.) including AttB1.1 and AttB2.1 cloning sites for Gateway™ cloning in the pDONR plasmid (Reece-Hoyes & Walhout, 2018). The synthetic transgene included a CaMV35S terminator sequence in 3’ position of the cystatin coding sequence and a CaMV35S promoter for constitutive expression in 5’ position (**Fig. 1A**).

### Transgenic lines

Transgenic potato lines were produced by *A. tumefaciens*-mediated transformation using 1-cm^2^ leaf explants from potato line K (*Solanum tuberosum*, cv. Kennebec) as source material (Badri et al., 2009). Transgenic cells were selected using the neomycin phosphotransferase II selection marker for kanamycin resistance and regenerated *in vitro* as described earlier (Michaud and Vrain, 1998). The regenerated plantlets were acclimated for 14 d in a growth chamber under a 24°C/21°C day/night temperature cycle, a 12:12 h light/dark photoperiod, a light intensity of 175 µmol.m^-2^.s^-1^ and a relative humidity of 60 %, before their transfer in greenhouse for multiplication and further analysis. Integration of the *nptii* selection transgene in kanamycin-resistant plants was confirmed by PCR using DNA extracted from the leaves of acclimated plants, according to Edwards *et al*. (1991). The following primers were used for amplification: 5’–ACTGAAGCGGGAAGGGACTGGCTGCTATTG and 3’–GATACCGTAAAGCACGAGGAAGCGGTCAG, to give a DNA amplicon visualized as a ∼500-base band after electrophoretic separation in 1% (w/v) agarose gels.

### Protein extraction and immunoblotting

Recombinant SlCYS8 was immunodetected in total soluble protein extracts from the third fully expanded leaf of ∼30 cm-tall plants, down from the apex. Leaf proteins were extracted in 50 mM Tris-HCl, pH 7.0, containing 1 mM sodium metabisulfite and 1 mM phenylmethylsulfonyl fluoride as described earlier (Rivard et al., 2006). The protein extracts were resolved by 12% (w/v) SDS-PAGE using a Bio-Rad Mini PROTEAN 3™ electrophoresis unit (Bio-Rad) and electrotransferred onto Hybond ECL nitrocellulose sheets (GE Healthcare) with a Mini-Trans-Blot Electrophoretic Transfer Cell™ (Bio-Rad), according to the supplier’s instructions. SlCYS8 was immunodetected with commissioned polyclonal IgG raised in rabbits against SlCYS8 (AgriSera) and goat anti-rabbit secondary IgG conjugated to alkaline phosphatase. Protein– antibody complexes were visualized using the alkaline phosphatase substrate 5-bromo-4-chloro-3-indolyl phosphate and nitro blue tetrazolium for color development (Life Technologies).

### Leaf proteome analysis

Leaf proteomes were analyzed by nanoscale liquid chromatography-tandem mass spectrometry (nanoLC-MS/MS) at the Plateforme protéomique du Centre hospitalier universitaire de Québec, Québec QC, Canada. Leaf protein samples from four biological replicates were analyzed for each tested line. Protein extracts (see above) were precipitated overnight at –20°C with five volumes of cold acetone, centrifuged and resuspended by sonication in 50 mM ammonium bicarbonate containing 1% (w/v) sodium deoxycholate. The resuspended proteins (50 µg) were reduced, alkylated and digested with sequence grade-trypsin (Promega) at a protease/protein ratio of 1:30. The resulting peptides were desalted using Empore StageTips pipette tips (CDS Analytical) prior to LC-MS/MS analysis. The nanoLC-MS/MS system comprised a U3000 NanoRSLC LC system (ThermoScientific Dionex Softron GmbH) in line with an Orbitrap Fusion Tribid-ETD mass spectrometer (ThermoScientific) driven by the Orbitrap Fusion Tune Application 3.3.2782.34 and equipped with a Nanospray Flex ion source. An equivalent of 2.0 µg protein of each sample diluted in 5 µL was injected into the system. The peptides were trapped for 5 min at 20 µL/min in loading solvent (2% ACN/0.05% TFA) on a 300mm i.d. x 5 mm, C18 PepMap100, 5 mm, 100 Å precolumn cartridge (Thermo Fisher Scientific). The precolumn was switched in line with a PepMap100 RSLC, C18 3 mm, 100 Å, 75 µm i.d. x 50 cm-long column (Thermo Fisher Scientific). The peptides were eluted along a 5%-40% linear gradient of solvent B (A: 0.1% FA; B: 80% ACN/0.1% FA) at 300 nL/min over a period of 90 min.

DATA-DEPENDENT ACQUISITION (DDA) Peptide spectra were acquired using the Thermo XCalibur software, v. 4.3.73.11, with lock mass on the *m/z* 445.12003 siloxane ion for internal calibration. Full-scan spectra (350-1800 *m/z*) were acquired in the Orbitrap using an AGC target of 4e5, a maximum injection time of 50 ms and a mass resolution of 120,000. Each MS scan was followed by the acquisition of fragmentation MS/MS spectra on the most intense ions for a total cycle time of 3 s (top speed mode). Selected ions were isolated using the quadrupole analyzer in a window of 1.6 m/z and fragmented by higher energy collision-induced dissociation with 35% collision energy. The resulting fragments were detected in the ion trap with a normalized AGC target of 33% and a maximal injection time of 50 ms. Dynamic exclusion of previously fragmented peptides was set to a period of 30 s and a tolerance of 10 ppm.

DATABASE SEARCHING Raw DDA data were searched against a UniProt Reference Proteome Database for *Solanum tuberosum* (ID UP000011115) using the MaxQuant software, with the “match between runs” function set and the following search parameters: ‘trypsin/P enzyme parameter’ enabled; carbamidomethylation of cysteine residues as a fixed modification; methionine oxidation and N-ter protein acetylation as variable modifications; mass search tolerances of 7 ppm and 0.5 Da for MS and MS/MS, respectively; and the peptide and protein identifications filtered at a False Discovery Rate of 1%. Proteins with a |z-score| ± 1.96 and a p-value < 0.05 were considered as differentially expressed in the assessed SlCYS8-expressing line compared to control line K.

### Hormone profile analysis

Hormones were quantified by high performance-liquid chromatography-electrospray ionization (HPLC-ESI)-MS/MS at the plant hormone profiling facility of the Canadian National Research Council Aquatic and Crop Resource Development Research Center, Saskatoon SK, Canada. Abscisic acid and catabolites, cytokinins, auxins and gibberellins were quantified as described by Lulsdorf et al. (2013). ACC was quantified according to Chavaux et al. (1997). The procedure for salicylic acid and jasmonic acid quantification was adapted from Murmu et al. (2014), and the procedure for salicylic acid hydrolysis of conjugated SA from Malamy et al. (1992). Analyses were monitored using the Multiple Reaction Monitoring (MRM) function of the MassLynx v. 4.1 control software (Waters). Resulting chromatographic traces are quantified off-line using the QuanLynx v4.1 software (Waters), wherein each trace is integrated and the resulting ratio of signals (nondeuterated/internal standard) compared with a previously constructed calibration curve to yield the amount of analyte present, in ng per sample. Calibration curves were generated with MRM signals obtained from standard solutions, based on the ratio of the chromatographic peak area for each analyte to that of the corresponding internal standard. QC samples, internal standard blanks and solvent blanks were prepared and analyzed along each batch of tissue samples.

### Osmotic stress experiment *in vitro*

The osmotic stress experiment was performed *in vitro* with control line K and SlCYS8-expressing lines C41, C43 and C51, using 15 plantlets per line. The plantlets were grown on solid Murashige and Skoog medium containing increasing concentrations of PEG (0 mM, 20 mM, 30 mM, 40 mM) to mimic drought conditions, according to the procedure of Anithakumari et al. (2011). The experimental setup followed a completely randomized factorial design, with plant lines and PEG concentrations as experimental variables. The plantlets were kept in glass tubes and let to grow for 28 d in a TCR60™ tissue culture room (Conviron, Winnipeg MB, Canada). Environmental conditions in the room were as follows: an ambient temperature of 20 °°C, a 16:8 h light/dark photoperiod, a light intensity of 175 µmol.m^-2^.s^-1^ and a relative humidity of 60%. Stem, leaf and root fresh and dry biomasses were measured at the end of the growing period. Root-*to*-shoot biomass ratios were calculated based on the dry biomass data.

### Water stress experiment in greenhouse

The greenhouse water deficit experiment was conducted with acclimated plants of control line K and SlCYS8-expressing lines C41, C43 and 51 at the High Performance Greenhouse Complex of Laval University (Québec QC, Canada), from September 2022 to January 2023. The experiment was arranged as a randomized complete block design with four repetitions of three soil matric potential (SMP) thresholds (–5 kPa, –10 kPa, –15 kPa). SMP thresholds were selected based on water retention curves, where SMP values and volumetric soil water data in the substrate were fitted to a PDI-variant of the bimodal unconstrained van Genuchten model. The soil substrate, constituted of two parts of sand (Saravia, #CH3386) for one part of organic soil (Fafard, SBG-55), was saturated with water before planting. Potato plants were grown in 0.14 m^3^ (L: 60 cm x H: 40 cm x W: 40 cm) plastic experimental units according to Matteau et al. (2021). For each experimental unit, two plants previously acclimated in the controlled environment chamber (see above) were planted at 7.5 cm under the soil surface. A tensiometer was installed at 10-cm depth in each experimental unit and tensiometer data were collected at two-min intervals. These measurements were used to calculate a mean SMP for each irrigation threshold to trigger automatic watering as needed, applied using drip irrigation emitters placed on the soil surface. Uniform irrigation across treatments was applied during the first 50 days following planting to ensure well-watered conditions during canopy growth. The water deficit experiment began at floral emergence and continued over 40 days. Canopy biomass was collected by cutting the stems off 15 cm above the soil. Tubers and roots were collected following senescence of the remaining plant parts for a period of 14 days.

### Tuber yield indices and statistical analyses

Analyses of variance (ANOVA) were computed using RStudio, v. 0.98.1103 to assess data generated *in vitro* for the PEG treatments and *ex vitro* with the greenhouse-grown acclimated plants. *Post-hoc* Tukey’s tests were performed for mean value comparisons on those ANOVA indicating (a) significant difference(s) among plant lines or stress treatments, using an alpha significance threshold of 5%. Drought tolerance indices (DTI) and yield performance indices (YTI) were calculated for total tuber biomass data as described in Ober et al. (2004), using the following formulas:

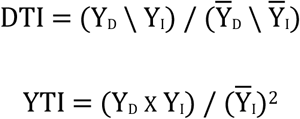

where Y_D_ is the tuber yield of a given genotype under a given water stress treatment, Y_I_ the tuber yield of this genotype in well-irrigated conditions, Y_D_ the average yield of all genotypes under a water stress treatment, and Y_I_ the average yield of all genotypes under well-irrigated conditions.

## Supporting information

Supplementary Dataset S1.xlxs

## Abbreviations

ANOVA: analysis of variance
CCII: corn cystatin II
DDA: data-dependent acquisition
DTI: drought tolerance index
LC-MS/MS: liquid chromatography-tandem mass spectrometry
PEG: polyethylene glycol
SlCYS8: eighth inhibitory domain of tomato multicystatin
YTI: yield performance index

## Acknowledgements

This work was supported by a Discovery grant from the Natural Science and Engineering Research Council of Canada to D.M.

## Supplementary material

**Supplementary Figure S1.** Optimization of the SlCYS8 codon sequence for heterologous expression in potato cells.

**Supplementary Figure S2.** Phytohormone contents in leaves of control line K and SlCYS8 lines C41, C43 and C51.

**Supplementary Table S1.** Complement to Fig. 3–Peptide ratios for defense/stress-related proteins up- or downregulated in SlCYS8-expressing lines C41, C43 and C51 compared to parent line K.

**Supplementary Table S2.** Growth parameters of *in vitro*-grown control K and SlCYS8-expressing potato plantlets under increasing concentrations of PEG used as an osmotic stress agent in the culture medium.

**Supplementary Table S3.** Complement to Table 1–Mean separation of growth parameter data from control line K and SlCYS8-expressing potato lines under well-irrigated or water stress regimes.

**Supplementary Table S4.** Complement to Fig. 6– Peptide ratios for primary metabolism-associated proteins up- or downregulated in SlCYS8-expressing lines C41, C43 and C51 compared to parent line K.

**Supplementary Dataset S1.** Raw and processed data from the LC-MS/MS analysis.

