## Supplementary Figures.pdf for "Tomato cystatin SlCYS8 as a trigger of drought tolerance and tuber yield in potato"

```
            1                                                50
  slcys8*   ATGAATCCTG GAGGAATTAC TAATGTACCT TTTCCTAATC TTCCTCAATT
   slcys8   ...AATCCTG GGGGCATTAC CAATGTTCCA TTCCCAAACC TCCCCGAGTT
Mutations   .......... .a..a..... t.....a..t ..t..t..t. .t..tc.a..

            51                                              100
  slcys8*   CAAGGACCTA GCTAGATTCG CAGTACAAGA TTACAACAAA AAGGAAAACG
   slcys8   CAAAGATCTT GCTCGTTTTG CTGTTCAAGA TTATAATAAG AAAGAGAATG
Mutations   ...g..c..a ...a.a..c. .a..a..... ...c..c..a ..g..a..c.

            101                                             150
  slcys8*   CACACCTTGA ATTCGTTGAG AACCTTAACG TTAAGGAACA GGTAGTAGCA
   slcys8   CTCATTTGGA GTTTGTAGAA AATTTGAATG TGAAGGAACA AGTTGTTGCT
Mutations   .a..cc.t.. a..c..t..g ..cc.t..c. .t........ g..a..a..a

            151                                             200
  slcys8*   GGTATTATTT ATTATATTAC TCTTGTTGCT ACTGATGCTG GTAAAAAGAA
   slcys8   GGAATAATAT ACTATATAAC ACTTGTGGCA ACTGATGCTG GAAAGAAGAA
Mutations   ..t..t..t. .t.....t.. t.....t..t .......... .t..a.....

            201                                             250
  slcys8*   GATTTACGAG ACAAAAATCC TTGTAAAAGG ATGGGAAAAT TTTAAGGAAG
   slcys8   AATATATGAG ACGAAGATTT TGGTGAAGGG ATGGGAGAAT TTCAAGGAAG
Mutations   g..t..c... ..a..a..cc .t..a..a.. ......a... ..t.......

            251                                    291
  slcys8*   TACAAGAGTT TAAACTAGTA GGAGACGCTA CTAAGCCTTA G
   slcys8   TTCAAGATTT CAAGCTTGTT GGTGATGCCA CTAAG..... .
Mutations   .a.....g.. t..a..a..a ..a..c..t. .......... .
```

**Supplementary Figure S1.** Optimization of the SlCYS8 codon sequence for heterologous expression in potato cells. Synonymous mutations were introduced *in silico* to minimize sequence identity with the potato multicystatin DNA sequence while keeping the original amino acid sequence unchanged. SlCYS8*, optimized sequence used for potato genetic transformation; Mutations, synonymous mutations introduced in the original *slcys8* transgene sequence.

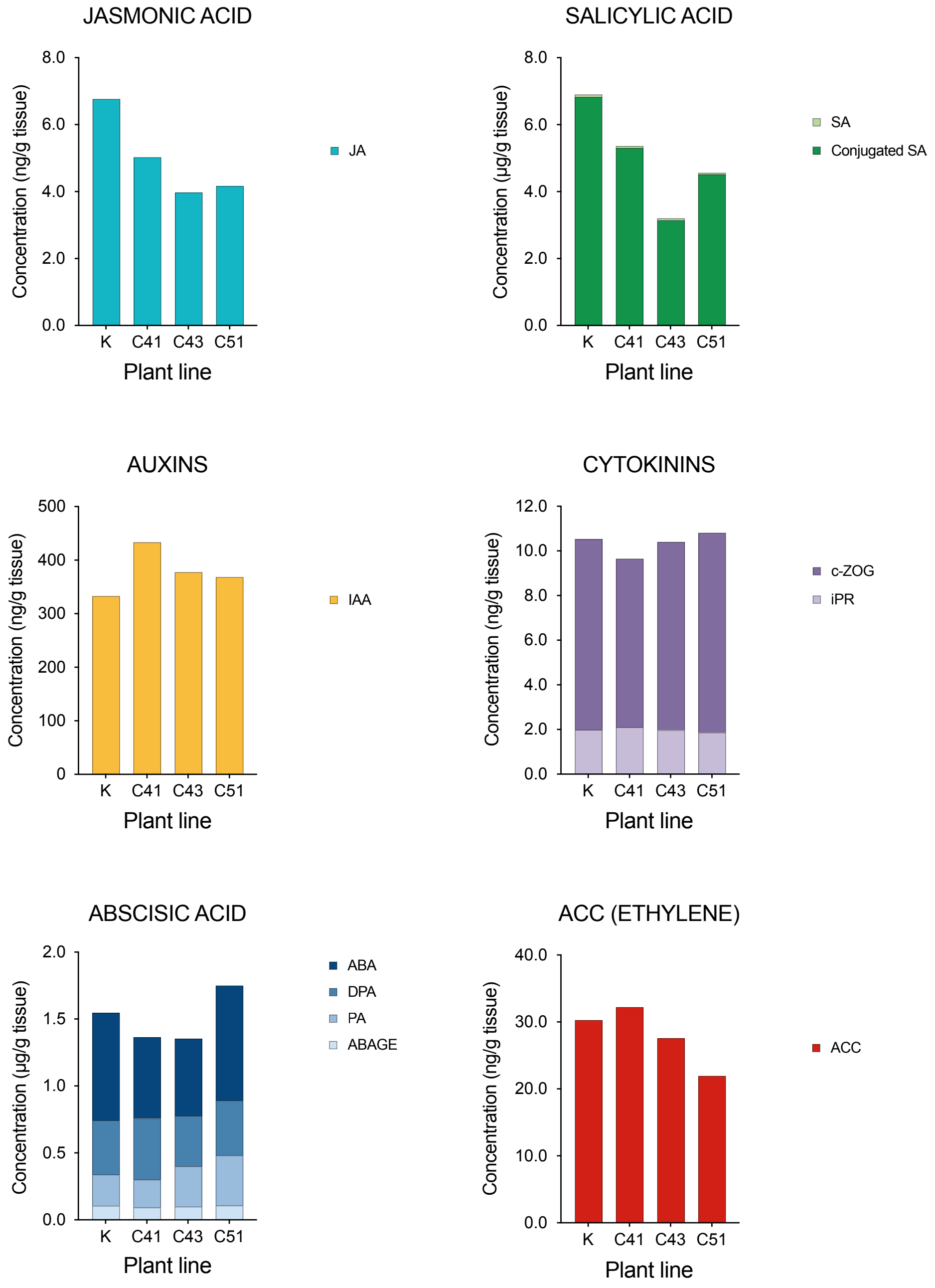


**Supplementary Figure S2.** Phytohormone contents in leaves of control line K and SlCYS8 lines C41, C43 and C51. ABA, *cis*-abscisic acid; ABAGE, abscisic acid glucose ester; ACC (ethylene direct precursor), 1-aminocyclopropane-1-carboxylic acid; *c*-ZOG, (*cis*) zeatin-O-glucoside; DPA, dihydrophaseic acid; IAA, indole 3-acetic acid; iPR, isopentenyladenine riboside; JA, jasmonic acid; PA, dihydrophaseic acid; SA, salicylic acid.
