## Supplementary Tables.pdf for "Tomato cystatin SlCYS8 as a trigger of drought tolerance and tuber yield in potato"

**Supplementary Table S1.** Complement to Fig. 3–Peptide ratios for defense/stress-related proteins up- or downregulated in SlCYS8-expressing lines C41, C43 and C51 compared to parent line K.

| Uniprot | SlCYS8 line [1] | | | Protein |
|---|---|---|---|---|
| | C41 | C43 | C51 | |
| M1BGS9 | 2,21 | | 3,09 | Coumarate ligase |
| M1BUC8 | 3,07 | 2,33 | 2,55 | Acetyl-CoA synthetase |
| M1CM72 | | 2 | 2,34 | 2-oxoglutarate-dependent dioxygenase |
| M1AED8 | 1,83 | 1,84 | 2,36 | ABC Transporter |
| M1CQL5 | 1,78 | 1,84 | 2,18 | Short-chain dehydrogenase/reductase 2 |
| M1CCL8 | | 2,6 | 0,71 | PRF (R1 homologue) |
| M1APJ0 | 1,64 | 1,27 | 2,02 | CMV 1a interacting protein |
| M1BWT8 | 1,43 | 1,18 | 1,82 | Rhamnose synthase |
| M1B8N1 | 1,64 | 1,21 | 1,51 | Serine/threonine-protein kinase bri1 |
| M1CEU1 | 1,15 | 0,88 | 2,26 | aarF dom. (BC1 complex kinase 3/ABC1 protein) |
| M1DBN7 | 1,45 | 1,06 | 1,68 | Glycosyoltransferase |
| M1CQA3 | 1,34 | 1,29 | 1,56 | Cytochrome P450-type monooxygenase |
| M1CTP6 | 1,5 | 1,31 | 1,36 | ACT DOMAIN-CONTAINING PROTEIN ACR11 |
| M1A4Q9 | | 0,67 | 2,06 | Peptidyl prolyl isomerase |
| M0ZS04 | | 0,76 | 1,8 | NMDA receptor-regulated protein |
| M1CVS3 | | 1,4 | 0,86 | PR-2 (ß-glucanase) |
| M1BGA0 | 1,17 | | 1,08 | Trehalose synthase |
| M0ZU79 | 1,1 | 0,2 | 1,34 | Coumarate ligase |

| | | | | |
|---|---|---|---|---|
| M0ZWD8 | 0,79 | 0,8 | 0,82 | Pre-pro-cysteine proteinase RD19 |
| M1AX69 | 1,69 | 0,39 | 0,15 | Protein phosphatase 2C |
| M1D4T3 | 0,18 | 1,74 | -0,06 | 2-Hydroxyisoflavanone dehydratase |
| M1AY17 | 0,2 | 1,14 | 0,47 | Peroxidase |
| M1A2A4 | 0,42 | 1,07 | 0,23 | Pathogenesis protein (PR)-1 |
| M1CCJ9 | -0,11 | 1,12 | -0,06 | Peroxidase |
| M1BDA2 | 1,16 | -1,14 | 0,75 | Glysosyltransferase (Protein CDI) |
| M1AN80 | 0,04 | 0,93 | -0,57 | Phytophthora-inhibited protease 1 (Cys protease) |
| M1BPP7 | -0,19 | 0,95 | -0,59 | Pathogen/wound-inducible antifungal prot CBP20 |
| Q5XUG9 | -0,35 | 1,14 | -1,36 | Thaumatin-like protein (PR-5) |
| M1AGK5 | -0,09 | 0,39 | -1,22 | Endochitinase |
| P52406 | -0,93 | 0,06 | -0,68 | Endochitinase 4 (Fragment) |
| M1AZ62 | -0,32 | -0,94 | -0,48 | Alpha-rhamnosidase |
| M1A703 | -1,03 | 0,66 | -1,54 | PRp27 (PR-17 protein) |
| M1APC8 | -0,56 | -0,02 | -1,34 | Acidic class II 1,3-beta-glucanase (PR-2) |
| M0ZMA8 | -0,71 | 0,84 | -2,16 | TSI-1 protein (PR protein) |
| M0ZMG2 | -0,35 | -0,25 | -1,45 | Class II chitinase |
| M1CY02 | -0,84 | -0,07 | -1,17 | Leucine-rich repeat protein (LRR receptor kinase) |
| M1AA90 | -0,78 | -0,12 | -1,29 | Secreted glycoprotein EP1 (mannose-binding lectin) |
| Q3HRZ0 | -0,2 | -2,05 | 0,03 | Cinnamoyl-CoA reductase |
| M1AMY2 | -1,41 | -1,52 | 0,53 | Kunitz proteinase inhibitor |
| M1BEU2 | -0,49 | 0,15 | -2,07 | Ser/Thr prot kinase (LRR recept. kinase) |
| P31212 | -1,14 | -0,85 | -0,43 | Threonine dehydratase biosynthetic (Fragment) |
| M1CX91 | -0,89 | 0,58 | -2,14 | PR-2 (ß-glucanase) |

| | | | | |
|---|---|---|---|---|
| M1D7L6 | -0,74 | -0,37 | -1,46 | Acid endochitinase |
| Q3YJS9 | -1,1 | -0,87 | -0,61 | Putative inactive patatin |
| M1AMY3 | -1,48 | -0,84 | -0,5 | Cysteine proteinase inhibitor 1 (Kunitz) |
| M1AMY4 | -1,32 | -1,26 | -0,53 | Cysteine protease inhibitor 1 (Kunitz) |
| M1B2K2 | -1,78 | -0,7 | -0,78 | Cys/Ser protease inhibitor (Kunitz) |
| M1B8U4 | -0,69 | -0,6 | -2,09 | Pathogenesis protein (PR)-1 |
| M0ZMG3 | -1,17 | -1,25 | -1,54 | Acidic 27 kDa endochitinase |
| M0ZIE0 | -1,62 | 0,16 | -2,58 | PR-2 (ß-glucanase) |
| M0ZTQ3 | -1,91 | -1,25 | -1,91 | ET-responsive proteinase inhibitor 1 |
| M1BHH1 | -2,91 | -1,83 | -2,67 | Trypsin inhibitor |
| M0ZU24 | -2,82 | -2,87 | -3,31 | Heat shock 70 kDa protein |

[1] Data are expressed as $\log_2$ values of peptide ratios compared to line K (relative $\log_2$ value of 0.00).

**Supplementary Table S2.** Growth parameters of *in vitro*-grown control K and SlCYS8-expressing potato plantlets under increasing concentrations of PEG used as an osmotic stress agent in the culture medium. Data were collected after 28 d. Each value is the mean of 4 biological replicate values ± SEM.

| [PEG] | Line | Stem length (cm) | Root length (cm) | Fresh shoot biomass (g) | Dry shoot biomass (mg) | Fresh root biomass (cm) | Dry root biomass (mg) |
|---|---|---|---|---|---|---|---|
| 0 mM | K | 9.29 ± 0.36 | 10.90 ± 0.48 | 1.13 ± 0.04 | 75.9 ± 2.89 | 0.44 ± 0.04 | 24.0 ± 1.45 |
| | C41 | 12.08 ± 0.33 | 10.58 ± 0.78 | 1.34 ± 0.03 | 92.7 ± 2.07 | 0.45 ± 0.03 | 22.0 ± 1.25 |
| | C43 | 11.99 ± 0.49 | 9.49 ± 0.56 | 1.34 ± 0.03 | 94.1 ± 3.17 | 0.37 ± 0.02 | 21.9 ± 0.99 |
| | C51 | 11.61 ± 0.32 | 9.18 ± 0.56 | 1.33 ± 0.03 | 93.4 ± 2.41 | 0.41 ± 0.03 | 24.8 ± 1.75 |
| 20 mM | K | 6.61 ± 0.50 | 11.08 ± 0.31 | 0.63 ± 0.07 | 64.3 ± 4.77 | 0.34 ± 0.03 | 23.1 ± 1.27 |
| | C41 | 6.08 ± 0.61 | 9.49 ± 0.70 | 0.59 ± 0.05 | 71.1 ± 3.96 | 0.46 ± 0.03 | 33.6 ± 2.51 |
| | C43 | 5.36 ± 0.43 | 8.65 ± 0.35 | 0.47 ± 0.02 | 62.0 ± 3.90 | 0.31 ± 0.02 | 27.8 ± 1.95 |
| | C51 | 6.99 ± 0.73 | 8.63 ± 0.49 | 0.58 ± 0.08 | 71.1 ± 4.77 | 0.41 ± 0.03 | 33.7 ± 1.73 |
| 30 mM | K | 5.64 ± 0.63 | 9.32 ± 0.60 | 0.47 ± 0.05 | 61.5 ± 6.09 | 0.27 ± 0.03 | 20.0 ± 2.06 |
| | C41 | 4.21 ± 0.43 | 9.13 ± 0.54 | 0.41 ± 0.03 | 52.7 ± 3.83 | 0.34 ± 0.03 | 22.2 ± 2.12 |
| | C43 | 3.71 ± 0.37 | 8.32 ± 0.33 | 0.48 ± 0.03 | 65.1 ± 4.29 | 0.18 ± 0.02 | 15.5 ± 1.68 |
| | C51 | 4.28 ± 0.60 | 8.19 ± 0.44 | 0.61 ± 0.04 | 84.8 ± 5.88 | 0.28 ± 0.03 | 24.4 ± 1.99 |
| 40 mM | K | 4.43 ± 0.44 | 8.64 ± 0.71 | 0.42 ± 0.04 | 62.1 ± 4.98 | 0.30 ± 0.03 | 22.8 ± 1.94 |
| | C41 | 4.41 ± 0.60 | 8.87 ± 0.65 | 0.34 ± 0.02 | 55.0 ± 3.53 | 0.27 ± 0.02 | 25.1 ± 2.21 |
| | C43 | 2.52 ± 0.27 | 7.58 ± 0.45 | 0.43 ± 0.03 | 63.6 ± 5.63 | 0.14 ± 0.02 | 14.3 ± 1.44 |
| | C51 | 3.06 ± 0.32 | 7.89 ± 0.42 | 0.39 ± 0.03 | 57.8 ± 3.75 | 0.24 ± 0.03 | 20.9 ± 2.08 |

**Supplementary Table S3.** Complement to Table 1–Mean separation of growth parameter data from control line K and SlCYS8-expressing potato lines under well-irrigated or water-deficient regimes. Values are the mean of 4 biological replicate values ± SEM.

| Statistical group | Treatment | Stem length (cm) | Stem number | Fresh shoot biomass (g) | Dry shoot biomass (g) | Root length (cm) | Dry root biomass (g) | Dry root-to-shoot ratio |
|---|---|---|---|---|---|---|---|---|
| Irrigation threshold | – 5 kPa | a | ab | a | a | n.s. | a | b |
| | – 10 kPa | a | b | b | a | n.s. | ab | b |
| | – 15 kPa | b | a | c | b | n.s. | b | a |
| Potato Line | K | a | b | n.s. | a | n.s. | n.s. | b |
| | C41 | b | a | n.s. | a | n.s. | n.s. | ab |
| | C43 | b | a | n.s. | a | n.s. | n.s. | ab |
| | C51 | b | a | n.s. | a | n.s. | n.s. | a |

In each column, different letters indicate significantly different mean values (Post-hoc Tukey's test; $\alpha = 0.05$). n.s., not significant.

**Supplementary Table S4.** Complement to Fig. 6–Peptide ratios for primary metabolism-associated proteins up- or downregulated in SlCYS8-expressing lines C41, C43 and C51 compared to parent line K.

| Uniprot | SlCYS8 line [1] | | | Protein |
|---|---|---|---|---|
| | C41 | C43 | C51 | |
| M1B8A5 | 3,44 | 4,29 | 3,01 | Plastid RNA-binding protein |
| M1BMW6 | 1,96 | | 2,15 | Ribosome biogenesis regulatory protein |
| M1BST4 | 1,71 | | 1,86 | 26S proteasome regulatory subunit S3 |
| M0ZVC8 | 1,59 | | 1,96 | RNA binding protein |
| M1CBP6 | | 1,38 | 1,61 | Nuclear proteasome inhibitor UBLCP1 |
| M0ZNI9 | 3,11 | -1,07 | 2,31 | RNA polymerase |
| M1AXT4 | 1,16 | 1,22 | 1,51 | Phospho-2-dehydro-3-deoxyheptonate aldolase |
| M1AEF9 | 1,45 | 0,91 | 1,49 | Transcription initiation factor IIB-2 |
| M1ATE5 | 0,96 | 1,15 | 1,7 | Hsp90 co-chaperone AHA1 |
| M0ZPC7 | 1,36 | 0,72 | 1,36 | DnaJ |
| M0ZUP2 | 1,28 | -0,26 | 2,4 | Zuotin |
| P54778 | 0,98 | 1,08 | 0,91 | 26S proteasome regulatory subunit 6B |
| M1AIE7 | 0,87 | 0,64 | 1,4 | Pentatricopeptide repeat-containing protein |
| M1BNH3 | 1,11 | 1,03 | 0,76 | Xylem serine proteinase 1 |
| M1B5V8 | 1,07 | 0,57 | 1,22 | Glutaminyl-tRNA synthetase |
| M1CLR3 | 0,63 | 0,6 | 1,63 | Translation factor GUF1 homolog |
| M1AEB0 | 0,82 | 0,59 | 1,42 | RUVb-like helicase |
| M1AI35 | -0,08 | 0,99 | 1,9 | ATP-dependent RNA helicase |
| M1BED0 | 1,06 | 0,78 | 0,88 | Poly-A binding protein |
| M1D118 | 0,95 | 0,19 | 1,52 | U5 small nuclear ribonucleoprotein component |

| | | | | |
|---|---|---|---|---|
| M1B4K0 | 0,65 | 0,81 | 1,1 | ATP-dependent Clp protease ATP-binding subunit clpA homolog CD4A |
| M1C1B5 | 0,73 | 1,01 | 0,66 | Eukaryotic translation factor |
| M1BK14 | 0,77 | 0,02 | 1,39 | Histidyl-tRNA synthetase |
| Q27S24 | 0,5 | 0,55 | 0,83 | ATP-dependent Clp protease proteolytic subunit |
| M1BGF6 | 0,39 | 0,33 | 0,92 | Chloroplast protease |
| M1B5I2 | 0,41 | 0,39 | 0,82 | ATP-dependent Clp protease ATP-binding subunit clpA homolog CD4B |
| M0ZXG5 | 0,36 | 0,35 | 0,71 | ATP-dependent Clp protease proteolytic subunit |
| M1BPT0 | 0,92 | 1,01 | -0,53 | NFU domain protein 4 |
| M0ZHY5 | -0,32 | 0,91 | -0,69 | Carboxypeptidase |
| M1AZL5 | -0,92 | 0,13 | -0,37 | Plastid-specific 30S ribosomal protein 3 |
| M1A9K8 | -0,44 | -0,44 | -0,56 | Protease degQ |
| M1BUM3 | -1,3 | -0,47 | -0,47 | Ribosomal protein L5 |
| M0ZRH9 | 2,89 | | 2,46 | Small GTPase Rab2 |
| M1BAR8 | 2,78 | 1,1 | 1,61 | Importin (EMB2734) |
| M1CSF3 | 1,08 | 0,95 | 1,63 | Co-atomer subunit beta |
| M1BSD0 | 1,26 | | 1,12 | Protein transport protein Sec24C |
| P30172 | 1,65 | 1,1 | 0,74 | Actin ACT-100 |
| M1AH97 | 0,73 | 0,87 | 1,53 | Short-chain dehydrogenase TIC32 |
| M1AW83 | 1,08 | 0,81 | 1,16 | Beta-adaptin |
| M1BAA4 | 1,07 | 0,44 | 0,94 | Protein transport protein SEC23 |
| M1C7M2 | -1,34 | -0,65 | 0,24 | Sorting nexin-4 |
| M0ZNQ6 | -1,24 | -0,16 | -0,47 | RAB6-interacting protein |
| P08454 | -1,14 | -0,93 | -0,58 | RAB6-interacting protein |
| M1A8E9 | 0,87 | 1,37 | 2,17 | ATAB2 |
| M1AD01 | 1,24 | 1,22 | 1,89 | Chlorophyl a-b binding protein |

| | | | | |
|---|---|---|---|---|
| M1AJ37 | 1 | | 1,8 | Tetratricopeptide repeat protein |
| M1CNC4 | 1,23 | 1,51 | | Ferredoxin |
| M1ACM0 | 1,23 | 1,43 | 1,1 | Dihydroflavonal-4-reductase |
| M0ZXL9 | 1,07 | 1,1 | 1,56 | Fibrillin / Plastid-lipid-assoc. protein 4 |
| M1ABC6 | 1,18 | 0,72 | 1,55 | PTAC14 |
| M0ZIF6 | 0,71 | 0,93 | 1,59 | Calcium homeostasis regulator CHoR1 |
| M1D586 | 1,12 | 0,78 | 1,26 | PLASTID-LIPID-ASSOCIATED PROTEIN 4 |
| M1APW9 | 0,48 | 0,48 | 1,71 | Pheophorbide A oxygenase |
| M1AFT5 | 0,77 | 0,43 | 1,36 | Chlorophyll a-b binding protein |
| M1C0J5 | 0,93 | 1,06 | 0,52 | Thioredoxin HCF164, chloroplastic |
| M1BUJ0 | 0,8 | 1,07 | 0,34 | Ferredoxin |
| M1CH70 | 2,29 | 0,73 | 1,65 | ß-amylase |
| M1BFJ3 | 1,1 | 1,61 | 1,94 | Galactokinase |
| P30924 | 0,99 | 1,26 | 2,15 | 1,4-alpha-glucan-branching enzyme |
| M1AKM2 | 0,73 | 0,79 | 2,12 | Xylulose kinase |
| M0ZJU5 | 1,3 | | 1,03 | AMP-activated protein kinase |
| M1D1P2 | 1,09 | 0,06 | 1,4 | O-glycosyl compounds hydrolase |
| M1AJ27 | -0,57 | -1,26 | 0,04 | Beta-galactosidase |

[1] Data are expressed as $log_2$ values of peptide ratios compared to line K (relative $log_2$ value of 0.00).
